# A NKp80-based identification strategy reveals that CD56^neg^ NK cells are not completely dysfunctional in health and disease

**DOI:** 10.1101/2020.02.12.945105

**Authors:** Ane Orrantia, Iñigo Terrén, Alicia Izquierdo-Lafuente, Juncal A. Alonso-Cabrera, Victor Sandá, Joana Vitallé, Santiago Moreno, María Tasias, Alasne Uranga, Carmen González, Juan J. Mateos, Juan C. García-Ruiz, Olatz Zenarruzabeitia, Francisco Borrego

**Affiliations:** Biocruces Bizkaia Health Research Institute, Immunopathology Group, 48903 Barakaldo, Spain.; Ramón y Cajal Health Research Institute (IRYCIS), Ramón y Cajal University Hospital, 28034 Madrid, Spain.; Hospital Universitari i Politecnic La Fe, 46026 Valencia, Spain.; Biodonostia Health Research Institute, Donostia University Hospital, 20014 Donostia-San Sebastián, Spain.; Biocruces Bizkaia Health Research Institute, Hematological Cancer Group, Cruces University Hospital, 48903 Barakaldo, Spain.; Ikerbasque, Basque Foundation for Science, 48013 Bilbao, Spain.

**Keywords:** CD16, CD56negative, HIV, NKp80, NK cells

## Abstract

Natural killer (NK) cells are usually identified by the absence of other lineage markers, due to the lack of a cell surface specific marker. CD56^neg^ NK cells, classically identified as CD56^neg^CD16^+^ are known to be expanded in some pathological conditions. However, studies on CD56^neg^ NK cells had revealed different results regarding the phenotype and functionality of these cells. This could be due to, among others, the unstable expression of CD16, which hinders CD56^neg^ NK cells identification. Hence, we aim to determine an alternative surface marker to CD16 to better identify CD56^neg^ NK cells. Using multiparametric flow cytometry, we have found that NKp80 is a good alternative to CD16 not only in healthy donors but also in HIV-1 infected subjects and multiple myeloma patients. Furthermore, we found differences between the functionality of CD56^neg^NKp80^+^ and CD56^neg^CD16^+^ NK cells both in healthy donors and patients, suggesting that the effector functions of CD56^neg^ NK cells are not as diminished as previously thought. We proposed NKp80 as a noteworthy marker to identify and accurately re-characterize human CD56^neg^ NK cells.

## INTRODUCTION

Natural killer (NK) cells constitute an essential part of the innate immune system, and they are able to eliminate virus-infected and tumor cells without previous sensitization[1, 2]. Based on the expression of CD56 and CD16, and the absence of CD3, three NK cell subsets can be distinguish: CD56^bright^CD16^+/-^, CD56^dim^CD16^+^ and CD56^neg^CD16^+^. The latter is very scarce in healthy donors, but it is expanded in chronic viral infections, such as HIV-1[3–6]. However, studies with similar cohorts of patients differ in the frequency, functionality and phenotype of the CD56^neg^ NK cell subset, which could be due to different CD56^neg^ NK cell identification strategies[7–10]. Furthermore, it is known that CD16 is downregulated by cryopreservation[11], after cytokine activation and target cell stimulation[12, 13]. Thus, the usage of this marker could lead to an inaccurate identification of CD56^neg^ NK cells and therefore inconsistent results.

NKp80 is an activating receptor expressed by virtually all fresh and activated mature NK cells[14]. NKp80 marks a critical step in NK cell development[15] and is a NK cell-specific marker among human innate lymphoid cells (ILCs)[16]. However, as far as we know, the possibility of using this marker to better identify CD56^neg^ NK cells has not been yet explored.

## RESULTS AND DISCUSSION

The CD16 receptor has traditionally been used, in combination with CD56, to identify the three major subsets of NK cells, with CD56^neg^ NK cells defined as CD56^neg^CD16^+^. However, CD16 is known to be downregulated in some situations, such as, cryopreservation, after cytokine activation and target cell stimulation [11–13]. With the aim to identify a more accurate marker with a more stable expression, we first compared CD16 with NKp80 receptor to identify CD56^neg^ NK cells in healthy donors (Fig EV1A). As expected, CD16 expression was downregulated in cryopreserved samples; however, the expression of NKp80 was not significantly altered after cell freezing (Fig EV2), suggesting that this receptor is more suitable for the detection of CD56^neg^ NK cells when it concerns to frozen cells. In addition, although no significant differences were seen regarding the percentage of CD56^neg^ NK cells selected using both markers (Fig EV1B), there was a higher frequency of Eomes^+^ cells in CD56^neg^NKp80^+^ subpopulation than in CD56^neg^CD16^+^ cells (Fig 1A). Eomes expression is frequently used as an intracellular marker for NK cells, given that it is a T-box transcription factor needed for the development and function of NK cells [16]. As the percentage of Eomes^+^ cells within CD56^neg^CD16^+^ cells was low, we considered the possibility that other CD16^+^ non-NK cells could have been selected using this gating strategy. This hypothesis was strengthened by the fact that within the CD56^neg^CD16^+^ population, Eomes^-^ cells had larger size than Eomes^+^ cells (Fig 1B). Thus, we studied the expression of CD123 receptor (α-chain of the interleukin 3 receptor) expressed, among others, in plasmacytoid dendritic cells (pDCs) [17] and monocytes, which are characterized by a larger size and granularity. Results showed that CD56^neg^CD16^+^Eomes^-^ cells expressed CD123, in contrast to CD56^neg^NKp80^+^Eomes^-^ cells that barely did (Fig 1C). Furthermore, the addition of an anti-CD123 mAb to the exclusion channel showed that the frequency of CD56^neg^CD16^+^Eomes^+^ cells significantly increased but still tended to be lower compared with CD56^neg^NKp80^+^Eomes^+^ cells (Fig 1D). This suggested that the inaccuracy in the identification of CD56^neg^ NK cells using CD16 is due to the selection of Eomes^-^ cells that, at least in part, could be pDCs or monocytes.

**Fig 1.**
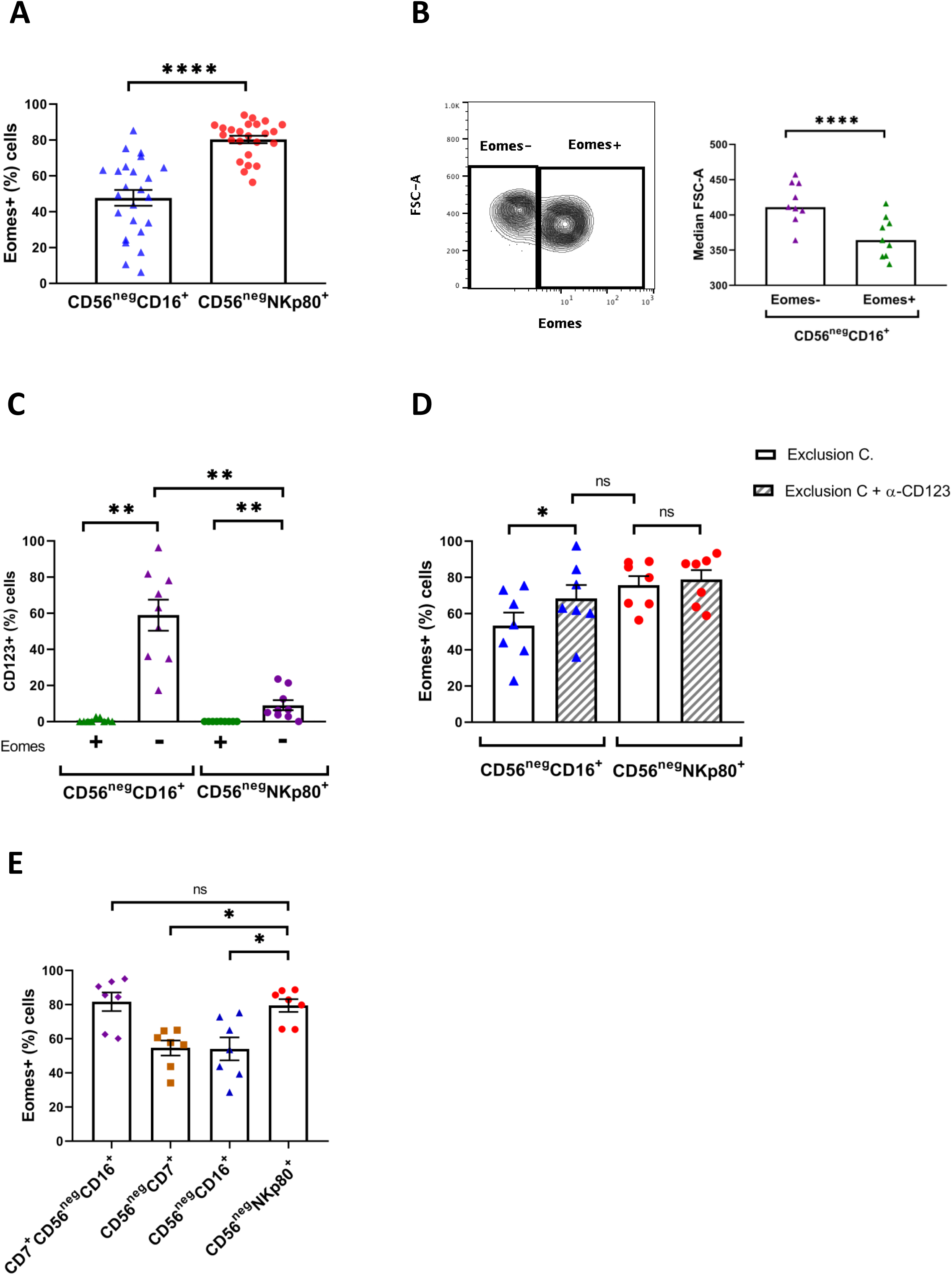
NKp80 better identifies CD56^neg^ NK cells than CD16 in healthy individuals. (A) Bar graph showing the percentage of Eomes^+^ cells within CD56^neg^CD16^+^ and CD56^neg^NKp80^+^ populations. (B) Left part, representative contour plot showing the Eomes expression and the size (FSC-A) of CD56^neg^CD16^+^ NK cells. Data from a representative healthy donor is shown. Numbers indicate the median of FSC-A. Right part, bar graph showing the median of FSC-A parameter within CD56^neg^CD16^+^Eomes- and CD56^neg^CD16^+^Eomes^+^ populations. (C) Bar graph showing the percentage of CD123^+^ cells within CD56^neg^CD16^+^Eomes^+^, CD56^neg^CD16^+^Eomes^-^, CD56^neg^NKp80^+^Eomes^+^ and CD56^neg^NKp80^+^Eomes^-^ populations. (D) Bar graph showing the percentage of Eomes^+^ cells within CD56^neg^CD16^+^ and CD56^neg^NKp80^+^ populations with or without the addition of anti-CD123 mAb to the exclusion channel (Exclusion C.). (E) Bar graph showing the percentage of Eomes^+^ cells within CD7^+^CD56^neg^CD16^+^, CD56^neg^CD7^+^, CD56^neg^CD16^+^ and CD56^neg^NKp80^+^ populations. The mean with the standard error of the mean (SEM) is represented, except for (B) in which the median is represented. Each dot represents a donor. *p<0.05, **p<0.01, **** p<0.0001, ns: not significant.

More recently, it was shown that including CD7 as an additional marker to study CD56^neg^CD16^+^ NK cells was an effective method to identify CD56^neg^ NK cells [10, 18]. Therefore, we compared the frequency of Eomes^+^ cells using CD7 or NKp80 markers to select CD56^neg^ NK cells. There was no significant difference between CD7^+^CD56^neg^CD16^+^ and CD56^neg^NKp80^+^ cells in terms of Eomes expression. However, using only CD7 instead of NKp80 showed that the frequency of Eomes^+^ cells in the CD7^+^CD56^neg^ population was much lower than in both CD7^+^CD56^neg^CD16^+^ and CD56^neg^NKp80^+^ cells (Fig1E). Nonetheless, combining CD7 and CD16 markers to identify the CD56^neg^ NK cell subset could be an obstacle in some situations, especially due to the unstable expression of the CD16 receptor and the need for an additional mAb and flow cytometer detector, which could be overcome by only using NKp80 as a marker.

Next, we studied CD300a [19–21] and 2B4 [22] (CD244) receptors that are expressed in NK cells, as markers for the identification of the CD56^neg^ NK cells. Results showed a lower frequency of Eomes+ cells both in CD56^neg^CD300a^+^ and in CD56^neg^2B4^+^ cells compared with CD56^neg^NKp80^+^ cells. Moreover, the addition of an anti-CD123 mAb to the exclusion channel did not increase the frequency of Eomes^+^ cells (Fig EV3).

As CD56^neg^ NK cells are infrequent in the peripheral blood of healthy donors, we next evaluated the accuracy of NKp80 to identify the expanded CD56^neg^ NK cells in pathological conditions (Fig EV4). No significant differences were noticed in the frequency of Eomes^+^ cells between CD56^neg^CD16^+^ and CD56^neg^NKp80^+^ subpopulations in untreated HIV-1 infected subjects. However, the frequency of Eomes^+^ cells was significantly higher in CD56^neg^NKp80^+^ than in CD56^neg^CD16^+^ cells in HIV-1 infected subjects under combined antiretroviral therapy (cART) (Fig 2A). The differences between patients groups could be explained because the relative frequency of CD56^neg^CD16^+^ non-NK cells (Eomes^-^) is lower in untreated patients due to a higher expansion of the CD56^neg^CD16^+^ NK cells (Eomes^+^) [5]. Thus, CD16 could only serve to identify CD56^neg^ NK cells in certain pathological conditions in which this subset is highly expanded. In addition, a higher frequency of Eomes^+^ cells within CD56^neg^NKp80^+^ NK cells was also noticeable in multiple myeloma patients (Fig 2B), in which CD56^neg^ NK cell expansion is more similar to the one of HIV-1 infected subjects under cART (Fig EV4). These findings suggest that NKp80, as demonstrated in healthy donors, is a noteworthy alternative to CD16 as a marker to identify CD56^neg^ NK cells also in disease.

**Fig 2.**
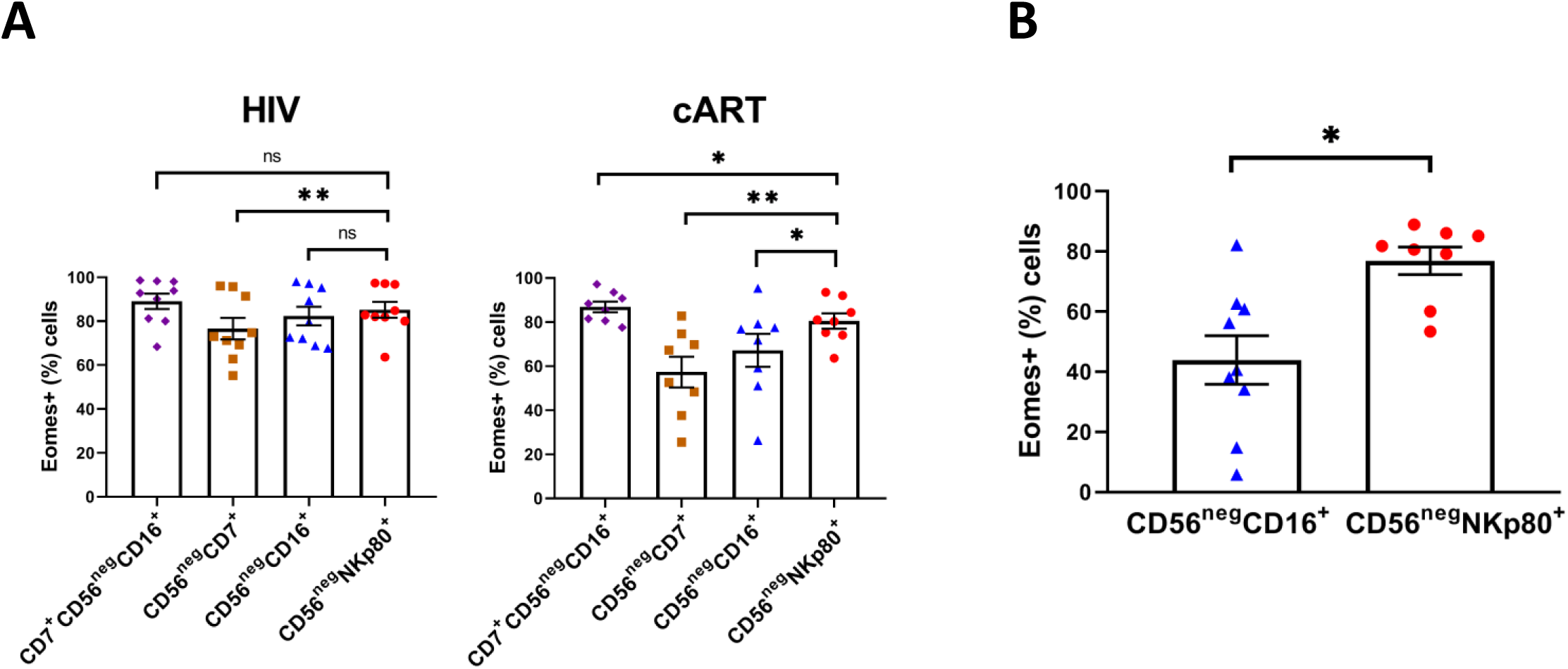
NKp80 better identifies CD56^neg^ NK cells than CD16 in pathological conditions. (A) Bar graphs showing the percentage of Eomes^+^ cells within CD7^+^CD56^neg^CD16^+^, CD56^neg^CD7^+^, CD56^neg^CD16^+^ and CD56^neg^NKp80^+^ populations in untreated HIV-1 infected subjects (HIV) and HIV-1 infected patients under cART (cART). (B) Bar graph showing the percentage of Eomes^+^ cells within CD56^neg^CD16^+^ and CD56^neg^NKp80^+^ populations in multiple myeloma patients. The mean with the standard error of the mean (SEM) is represented. Each dot represents a donor. *p<0.05, **p<0.01, ns: not significant.

CD56^neg^ NK cells have been described as functionally impaired compared to CD56^dim^ NK cells [7,8,10,23]. However, others have proposed that these cells are skewed rather than dysfunctional [9]. These differences may be due to inaccurate identification of CD56^neg^ NK cells using CD16. Therefore, we studied the effector functions of CD56^neg^CD16^+^ and CD56^neg^NKp80^+^ cells from healthy donors and HIV-1 infected subjects, by measuring degranulation (CD107a) [24] and production of TNF and IFNγ after cytokines and K562 target cell stimulations [25]. Results showed that NKp80 was not significantly downregulated after K562 cell line and cytokine stimulation, while CD16 expression decreased after stimulation (Fig EV5). CD56^neg^NKp80^+^ cells exhibited higher production of TNF and IFNγ than CD56^neg^CD16^+^ cells in untreated and treated HIV-1 infected subjects (Fig 3A). Furthermore, in healthy donors, both cytokine production and the degranulation capability were increased in the CD56^neg^NKp80^+^ cells (Fig 3B). Our results indicate that, although CD56^neg^ NK cells have lower effector functions than CD56^dim^ NK cells (Fig EV6), their functionality might not be completely impaired [26].

**Fig 3.**
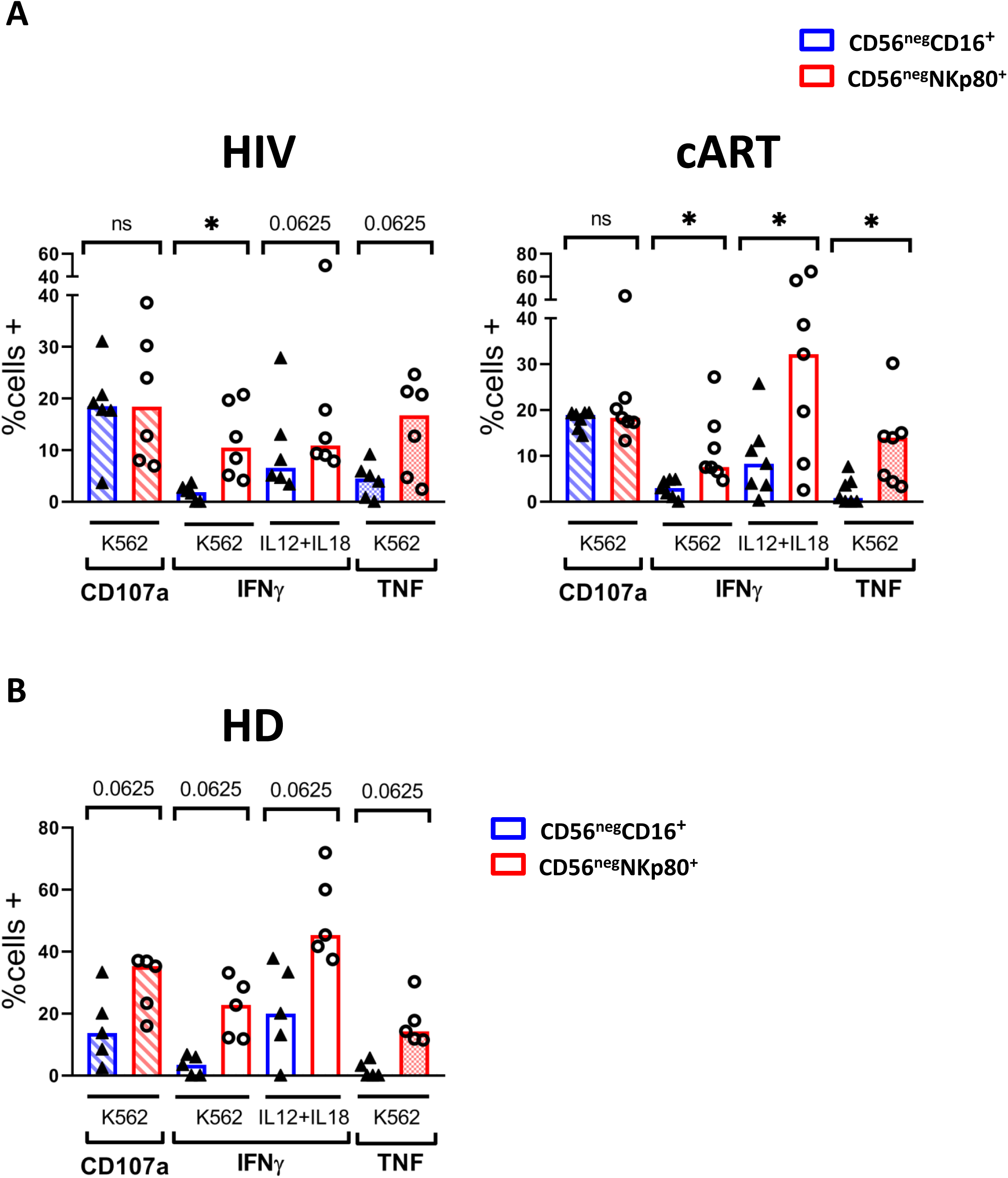
Degranulation (CD107a) and cytokine production (IFNγ and TNF) by CD56^neg^ NK cells in response to K562 cell line and IL-12+IL-18 stimulation. (A) Bar graphs showing the percentage of positive cells for CD107a, IFNγ and TNF from HIV-1 infected subjects (HIV) and HIV-1 infected patients under cART (cART) after stimulation with the K562 cell line and IL-12+IL-18 cytokines within CD56^neg^CD16^+^ and CD56^neg^NKp80^+^ NK cells. (B) Bar graph showing the percentage of positive cells for CD107a, IFNγ and TNF from healthy donors (HD) after stimulation with the K562 cell line and IL-12+IL-18 cells within CD56^neg^CD16^+^ and CD56^neg^NKp80^+^ NK cells. The median is represented. Each dot represents a donor. *p<0.05, ns: not significant.

In conclusion, in this study we have demonstrated that NKp80 is a more precise marker than CD16 in order to identify CD56^neg^ NK cells and that it is not downregulated after sample cryopreservation or cell activation. Importantly, using the NKp80 marker for the identification, we have demonstrated that the effector functions of CD56^neg^ NK cells are not as diminished as previously thought. Thus, an NKp80-based strategy for the better identification and re-characterization of this NK cell subset could help to clarify its function and relevance in health and disease.

## MATERIALS AND METHODS

### Subjects and samples

For this study, buffy coats from 24 healthy adult donors and cryopreserved peripheral blood mononuclear cells (PBMCs) from 9 multiple myeloma patients were collected through the Basque Biobank for Research (http://www.biobancovasco.org), which complies with the quality management, traceability and biosecurity, set out in the Spanish Law 14/2007 of Biomedical Research and in the Royal Decree 1716/2011. The study was approved by the Basque Ethics Committee for Clinical Research (PI2014017 and PI+CES+INC-BIOEF 2017-03). All subjects provided written and signed informed consent in accordance with the Declaration of Helsinki. In addition, cryopreserved PBMCs from healthy donors (n=5), untreated HIV-1 infected subjects (n=9) and patients under cART (n=8) were kindly provided by the HIV BioBank integrated in the Spanish AIDS Research Network (RIS) (Appendix I). Samples were processed following current procedures and frozen immediately after their reception. All patients participating in the study gave their informed consent and protocols were approved by institutional ethical committees.

All HIV-1 infected patients were asymptomatic when the sample was collected, were not co-infected with hepatitis C virus (HCV), had more than 200 CD4+ T cells/mm^3^ and they had never been diagnosed with AIDS. Untreated HIV-1 infected subjects had detectable viremia (>10,000 HIV-RNA copies/ml) and they had never been treated with cART, while patients under cART had undetectable viremia and had been treated with cART at least for 6 months. Clinical data of HIV-1 infected patients were obtained from the RIS database. Clinical data are shown in Supplementary Table 1.

### Antibodies and reagents

For flow cytometry-based procedures, the following fluorochrome-conjugated anti-human monoclonal antibodies (mAbs) were used: Brilliant Violet (BV)421 anti-CD56 (NCAM 16.2), BV510 anti-CD3 (UCHT1), BV510 anti-CD14 (MФP9), BV510 anti-CD19 (SJ25C1), BV510 anti-CD123 (9F5), PE anti-CD123 (9F5), PE anti-CD7 (M-T701) and PerCP-Cy5.5 anti-IFNγ (B27) from BD Biosciences; FITC anti-CD16 (B73.1) and APC anti-TNF (MAb11) from BioLegend; PE anti-CD300a (E59.126) and PE anti-2B4 (clone C1.7) from Beckman Coulter; PE anti-CD107a (REA792) and PE-Vio770 anti-NKp80 (4A4.D10) from Miltenyi Biotec; eFluor660 anti-Eomes (WD1928) from eBioscience. Dead cells were detected with the LIVE/DEAD™ Fixable Aqua Dead Cell Stain Kit for 405nm excitation from Invitrogen, following manufacturer’s protocol.

The following reagents were also used: Foxp3/Transcription Factor Staining Buffer Set from eBioscience; Brilliant Stain Buffer, BD GolgiStop™ Protein Transport Inhibitor (monensin), BD GolgiPlug™ Protein Transport Inhibitor (brefeldin A) and BD Perm/Wash™ Buffer from BD Bioscience; and paraformaldehyde (PFA) from Sigma-Aldrich/Merck.

### Flow cytometry

Fresh PBMCs from healthy donors were obtained from buffy coats by Ficoll (GE Healthcare) density gradient centrifugation and cryopreserved in Fetal Bovine Serum (FBS) (GE Healthcare Hyclone) with 10% Dimethylsulfoxide (DMSO) (Thermo Scientific Scientific).

For phenotypical studies, cryopreserved PBMCs from healthy donors, HIV-1-infected subjects and multiple myeloma patients were thawed at 37°C and washed twice with RPMI 1640 medium with L-Glutamine (Lonza). Then, cells were incubated for 1 hour at 37°C with 10U DNase (Roche) in R10 medium (RPMI 1640 medium containing GlutaMAX from Thermo Fisher Scientific, 10% FBS and 1% Penicillin-Streptomycin from Thermo Fisher Scientific). Afterwards, cells were counted and washed with Phosphate Buffered Saline (PBS) (Gibco, Thermo Fisher Scientific). Then, dead cells were excluded by using the LIVE/DEAD reagent. For the staining of NK cell surface markers, cells were first washed with PBS containing 2.5% of Bovine Serum Albumin (BSA) (Millipore) and then incubated for 30 minutes at 4°C with fluorochrome-conjugated mAbs. To identify NK cells, first viable cells that were negative for CD3, CD14 and CD19 were electronically gated, and then, by using the anti-CD56 mAb in combination with mAbs against CD16, NKp80, CD300a, 2B4 and/or CD7 NK cells were classified in three subsets: CD56^bright^, CD56^dim^ and CD56^neg^. After this, cells were washed again with 2.5% BSA in PBS and fixed and permeabilized with Foxp3/Transcription Factor Staining Buffer Set following manufacturer’s recommendations. Finally, cells were stained using anti-Eomes mAb for 30 minutes at room temperature (RT) and washed with Permeabilization Buffer 1x (eBioscience). Sample acquisition was carried out in a MACSQuant Analyzer 10 flow cytometer (Miltenyi Biotec).

For functional assays, after DNase treatment, PBMCs from HIV-1-infected subjects and healthy donors were counted and plated at 0.5 x 10^6^ cell/well in 48 well plates in NK cell culture medium (RPMI 1640 medium with GlutaMAX, 10% FBS, 1% penicillin-streptomycin, 1% non-essential amino acids and 1% Sodium-Pyruvate). PBMCs were then primed with interleukin (IL)-15 (10ng/mL) and cultured overnight (ON). For cytokine stimulation, IL-12 (10ng/mL) and IL-18 (50ng/mL) were also added to plated PBMCs before the ON culture. For target cell stimulation, K562 cells were added after ON culture at Effector:Target (E:T) 1:1 ratio (0.5×10^6^ PBMCs and 0.5×10^6^ K562 cells) and cells were incubated for 6h. CD107a was added at the start of the coculturing period and protein transport inhibitors were added after 1 hour for the rest of the incubation time following manufacturer’s protocol. Afterwards, viability and surface marker staining was performed as explained above. For intracellular staining, cells were fixed with 4% PFA for 15 minutes on ice and then washed twice with 2.5% BSA in PBS. After this, cells were permeabilized with BD Perm/Wash Buffer 1X for 15 minutes at RT. Finally, the corresponding mAbs were added for 30 minute and cells were washed with BD Perm/Wash Buffer 1X before acquisition in the MACSQuant Analyzer 10 flow cytometer (Miltenyi Biotec). The percentage of positive cells for CD107a, IFNγ and TNF was calculated after subtracting the non-stimulus condition.

### Statistical analysis and data representation

Data were analysed using FlowJo™ Version 10.4.1. GraphPad Prism v8.01 software was used for graphical representation and statistical analysis. Data were represented showing means ± standard error of the mean (SEM) or median as indicated in the figure legend. Prior to statistical analyses, data were tested for normal distribution with Kolmogórov-Smirnov normality test. In the case of multiple myeloma patients, an outlier was identified and removed using Grubb test (alpha=0.05). If data were normally distributed, t test for paired values was used to determine significant differences. Non-normal distributed data were compared with Wilcoxon matched-pairs signed rank test. Kruskal-Wallis test was used for multiple comparisons of non-normal data (Fig EV4). *p<0.05, **p<0.01, ***p<0.001, ****p<0.0001.

## ACKNOWLEDGMENTS

The authors thank the healthy donors and patients who participated in the study and the staff from Basque Biobank for Research, the HIV BioBank, the Basque Center for Blood Transfusion and Tissues, Cruces University Hospital and Donostia University Hospital.

This study was supported by grants from AECC-Spanish Association Against Cancer (PROYE16074BORR), “Plan Estatal de I+D+I 2013–2016, ISCIII181 Subdirección de Evaluación y Fomento de la Investigación-Fondo Europeo de Desarrollo Regional (FEDER) (Grant PI13/00889)” and Marie Curie Actions, Career Integration Grant, European Commission (Grant CIG 631674). J.V. and I.T. are recipients of a predoctoral contract funded by the Department of Education, Basque Government (PRE_2018_2_0242 and PRE_2019_2_0109). I.T. is the recipient of a fellowship from the Jesús de Gangoiti Barrera Foundation (FJGB18/002). O.Z. is the recipient of a postdoctoral contract funded by ISCIII-Contratos Sara Borrell (CD17/0128) and the European Social Fund (ESF)-The ESF invests in your future. F.B. is an Ikerbasque Research Professor, Ikerbasque, Basque Foundation for Science. We want to particularly acknowledge the patients in this study for their participation and to the HIV BioBank integrated in the Spanish AIDS Research Network and collaborating Centers for the generous gifts of clinical samples used in this work. The HIV BioBank, integrated in the Spanish AIDS Research Network, is supported by “Instituto de Salud Carlos III”, Spanish Health Ministry (Grant n° RD06/0006/0035, RD12/0017/0037 and RD16/0025/0019) as part of the “Plan Nacional R+D+I” and co-financed by “ISCIII-Subdirección General de Evaluación y el Fondo Europeo de Desarrollo Regional (FEDER)”. This study would not have been possible without the collaboration of all the patients, medical and nursery staff and data managers who have taken part in the project. The RIS Cohort (CoRIS) is funded by the “Instituto de Salud Carlos III” through the “Red Temática de Investigación Cooperativa en SIDA” (RIS C03/173, RD12/0017/0018 and RD16/0002/0006) as part of the “Plan Nacional R+D+I” and co-financed by “ISCIII-Subdirección General de Evaluación y el Fondo Europeo de Desarrollo Regional (FEDER)”.

## AUTHOR CONTRIBUTIONS

A.O. designed and performed the experiments, analyzed and interpreted the data, designed the figures, and wrote the manuscript. A.I-L, J.A.A-C. and V.S. performed experiments. S.M., M.T., A.U., C.G., J.J.M. and J.C.G-R. clinically characterized the patients and participated in the interpretation of the data. J.V. and I.T. participated in the interpretation of the data. O.Z. participated in the design of the study and interpreted the data. F.B. conceived and designed the study, interpreted the data, and wrote the manuscript. All the authors critically reviewed, edited, and approved the final manuscript.

---

**Fig EV1.**
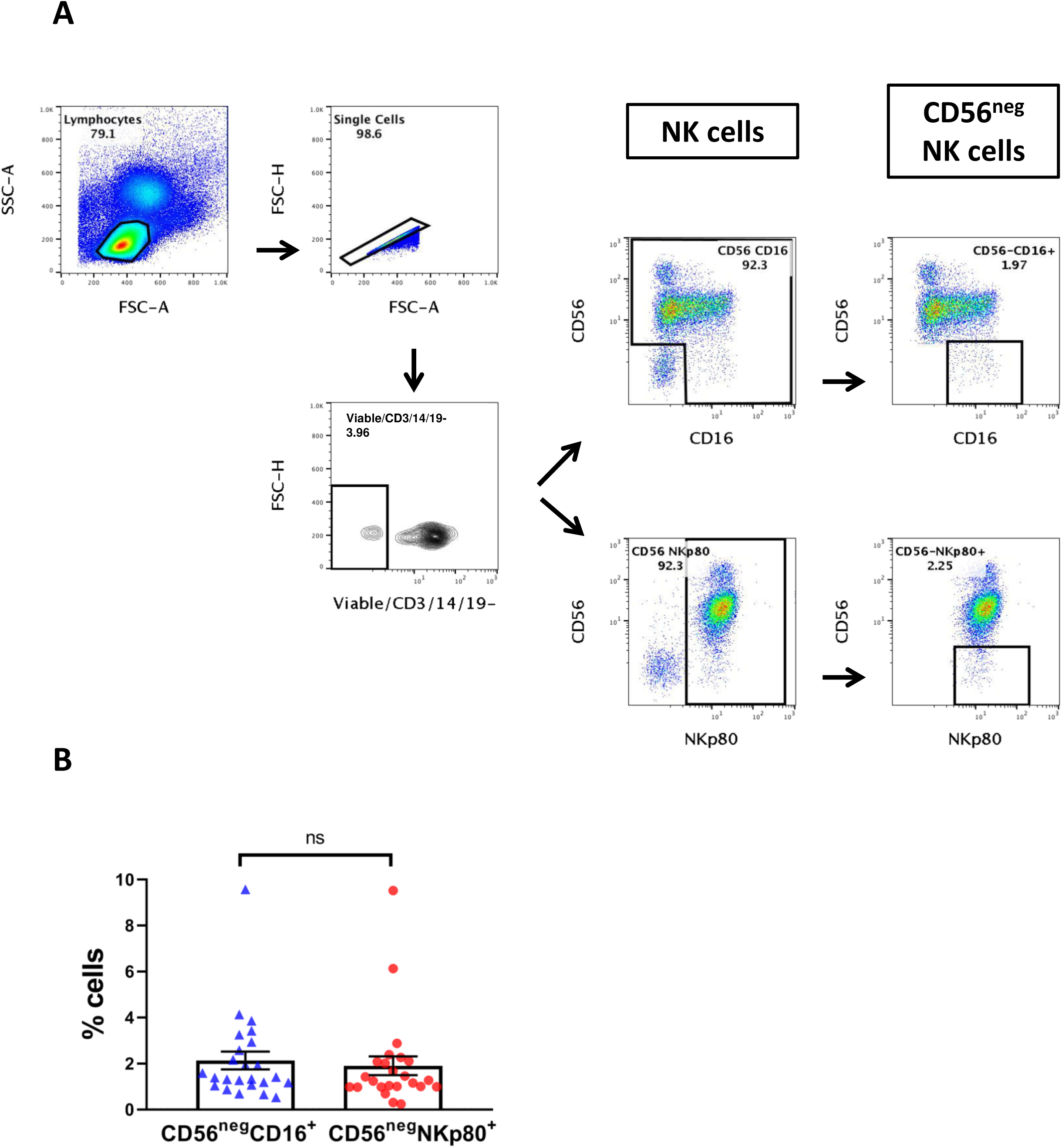
Identification of CD56^neg^ NK cells. (A). Pseudocolor and contour plot graphs representing the gating strategy utilized for the identification of CD56^neg^ NK cells. Data from a representative cryopreserved sample from a healthy donor is shown. Lymphocytes were electronically gated based on their forward and side scatter parameters and then single cells were selected. To identify NK cells, the population negative for the exclusion channel (viability, CD3, CD14 and CD19) was selected. Then CD56^neg^ NK cells were identified using different gating strategies. (B) Percentage of CD56^neg^CD16^+^ and CD56^neg^NKp80^+^ cells in healthy donors. Bar graph showing the percentage of CD56^neg^CD16^+^ and CD56^neg^NKp80^+^ cells in healthy donors. The mean with the standard error of the mean (SEM) is represented. Each dot represents a donor. ns: not significant.

**Fig EV2.**
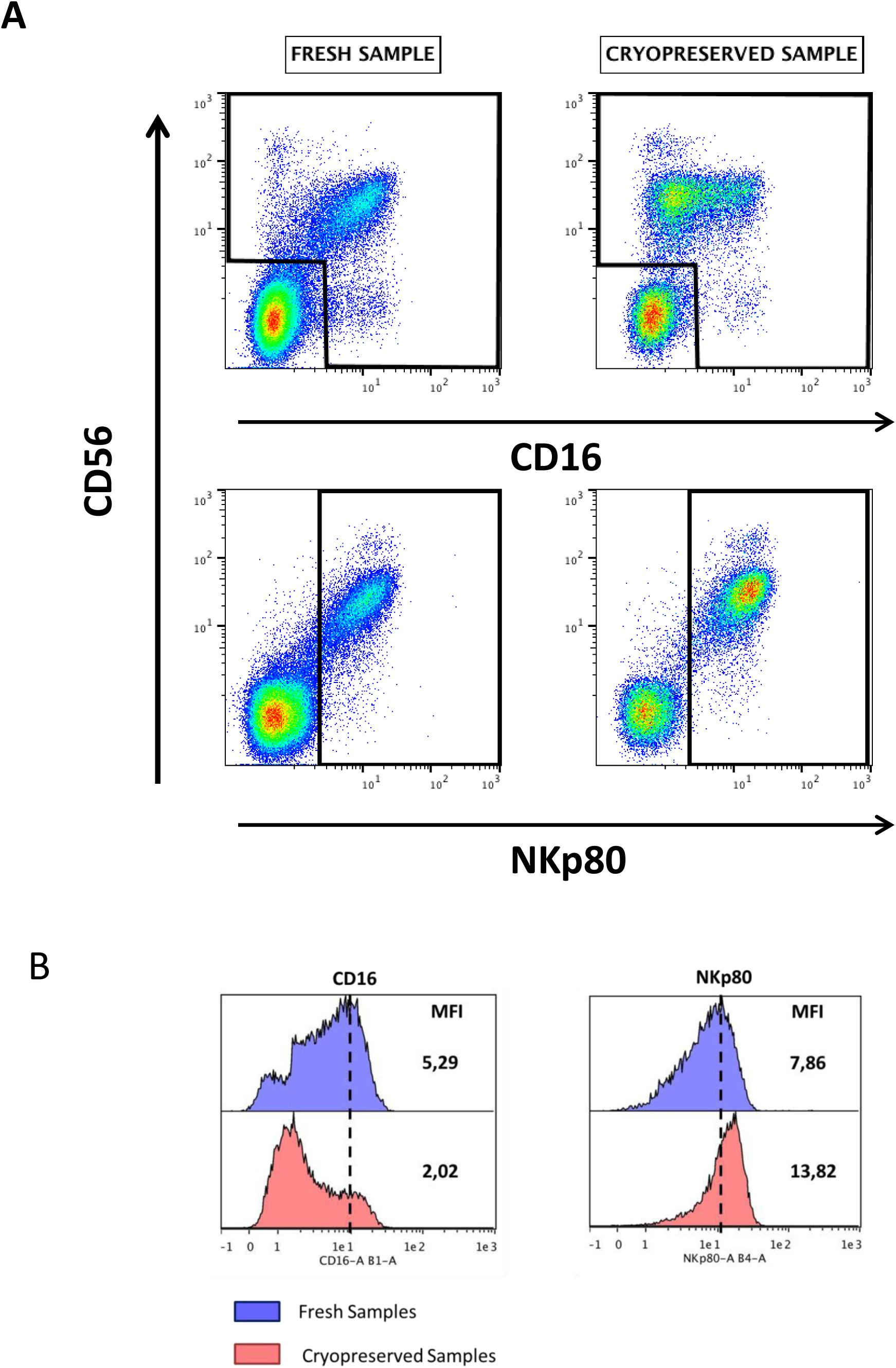
**CD16 but not NKp80 is downregulated after cell cryopreservation**. (A) Representative pseudocolor plot graphs comparing the expression of CD16 and NKp80 in fresh and cryopreserved samples. Data from a representative healthy donor is shown. (B) Histograms showing the median fluorescence intensity (MFI) of CD16 and NKp80 on NK cells in fresh and cryopreserved samples. Data from a representative healthy donor is shown.

**Fig EV3.**
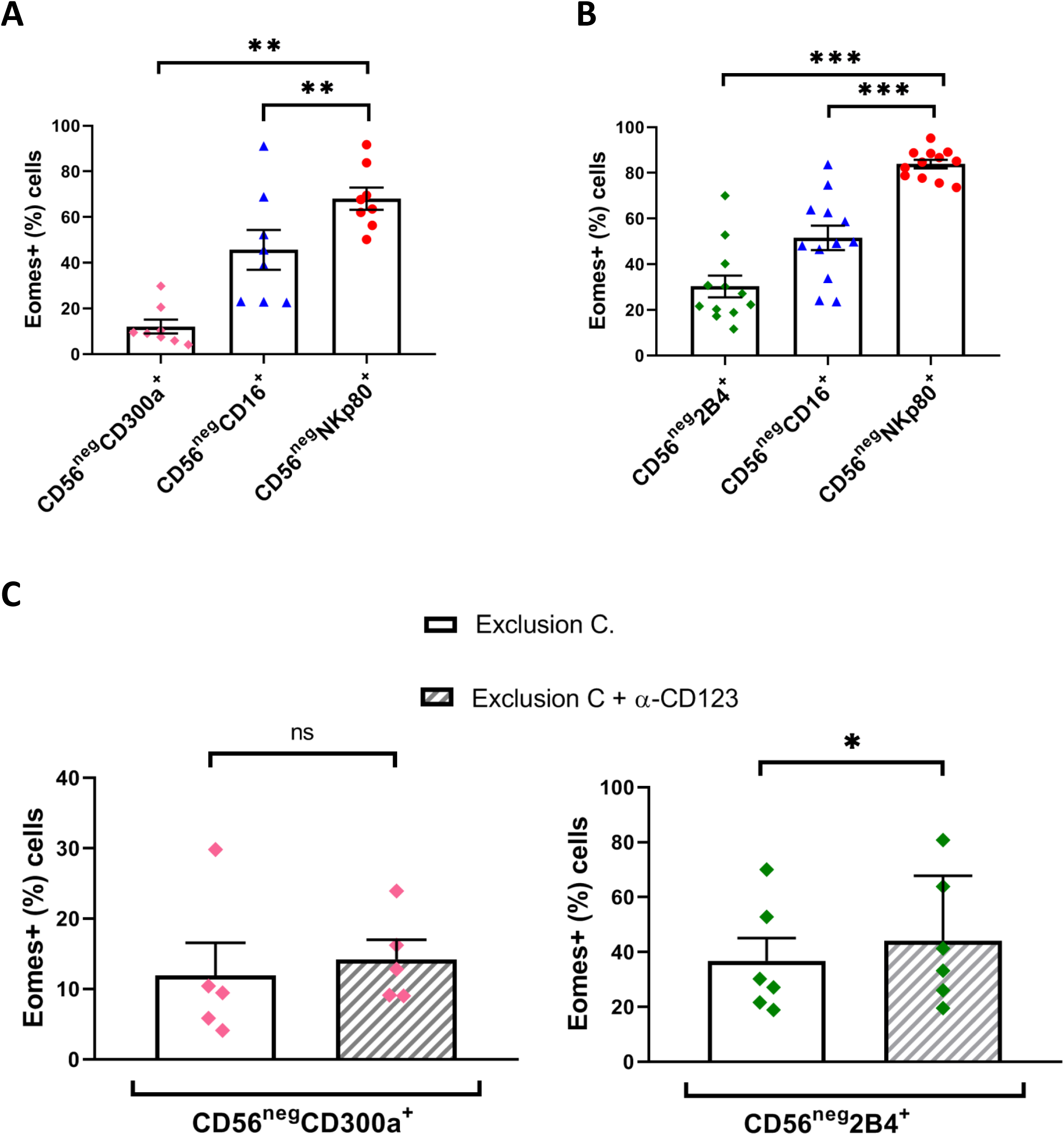
CD300a and 2B4 (CD244) receptors do not properly identify CD56^neg^ NK cells. (A) Bar graph showing the percentage of Eomes^+^ cells within CD56^neg^CD300a^+^, CD56^neg^CD16^+^ and CD56^neg^NKp80^+^ populations. (B) Bar graph showing the percentage of Eomes^+^ cells within CD56^neg^2B4^+^, CD56^neg^CD16^+^ and CD56^neg^NKp80^+^ populations. (C) Bar graphs showing the percentage of Eomes+ cells within CD56^neg^CD300a^+^ and CD56^neg^2B4^+^ populations with or without the addition of anti-CD123 mAb to the exclusion channel (Exclusion C.). The mean with the standard error of the mean (SEM) is represented. Each dot represents a donor. *p<0.05, **p<0.01, ***p<0.001, ns: not significant.

**Fig EV4.**
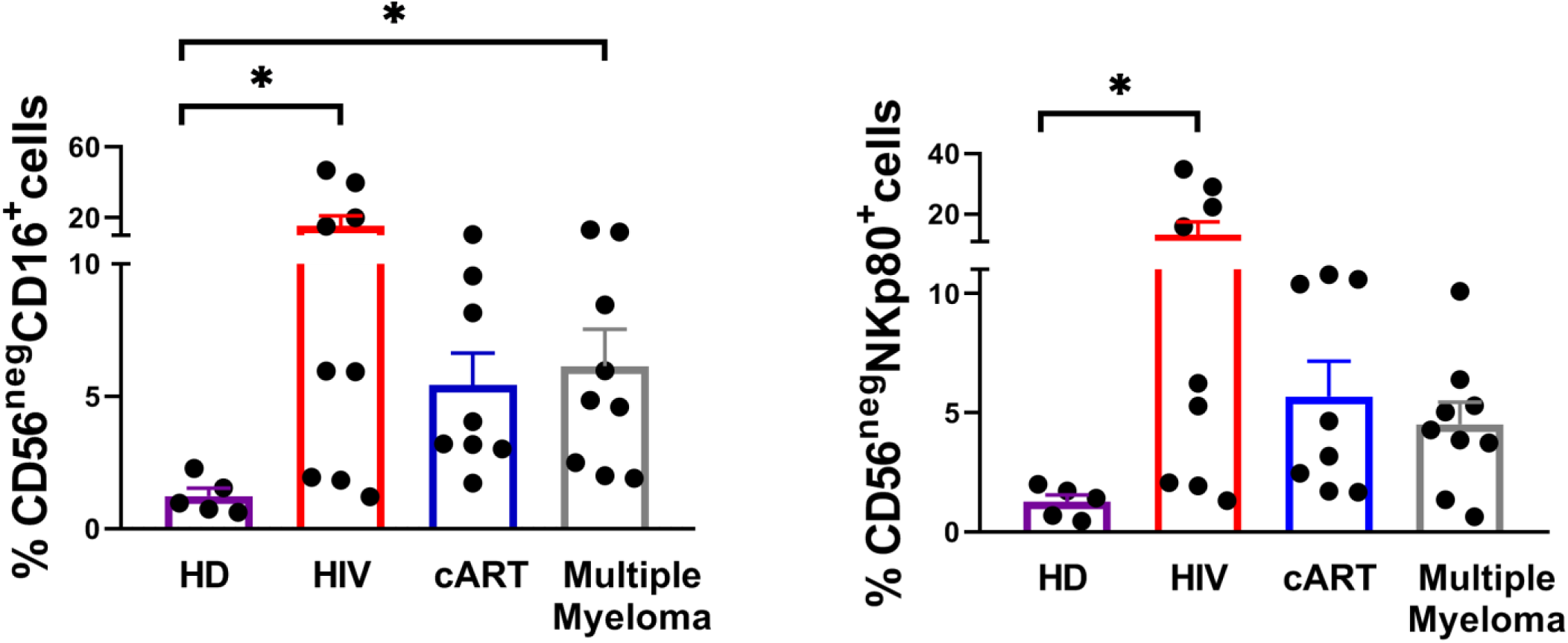
CD56^neg^ NK cells are expanded in pathological conditions. Bar graph showing the percentage of CD56^neg^CD16^+^ (left part) and CD56^neg^NKp80^+^ (right part) NK cells within healthy donors (HD), untreated HIV-1 infected subjects (HIV), HIV-1 infected patients under cART (cART) and multiple myeloma patients. The mean with the standard error of the mean (SEM) is represented. Each dot represents a donor. *p<0.05.

**Fig EV5.**
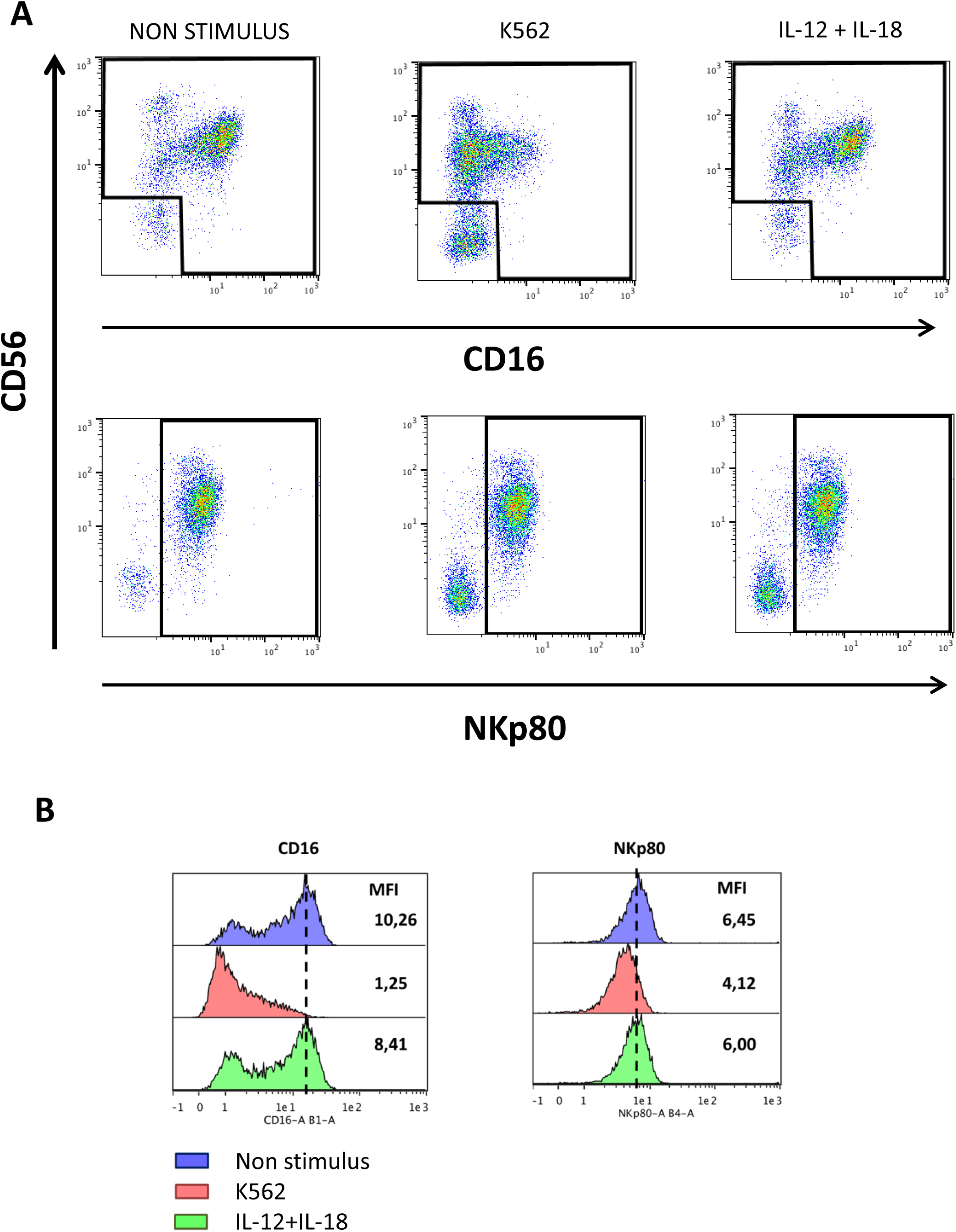
CD16, but no NKp80, is downregulated after K562 cell line and IL-12+IL18 cytokine stimulation. (A) Representative pseudocolor plot graphs comparing the expression of CD16 and NKp80 in non-stimulated condition, and after K562 cell line and IL-12+IL-18 cytokine stimulation. Data from a representative healthy donor is shown. (B) Histograms showing the median fluorescence intensity (MFI) of CD16 and NKp80 on NK cells in in non-stimulus condition and after K562 cell line and IL-12+IL-18 cytokine stimulation. Data from a representative healthy donor is shown.

**Fig EV6.**
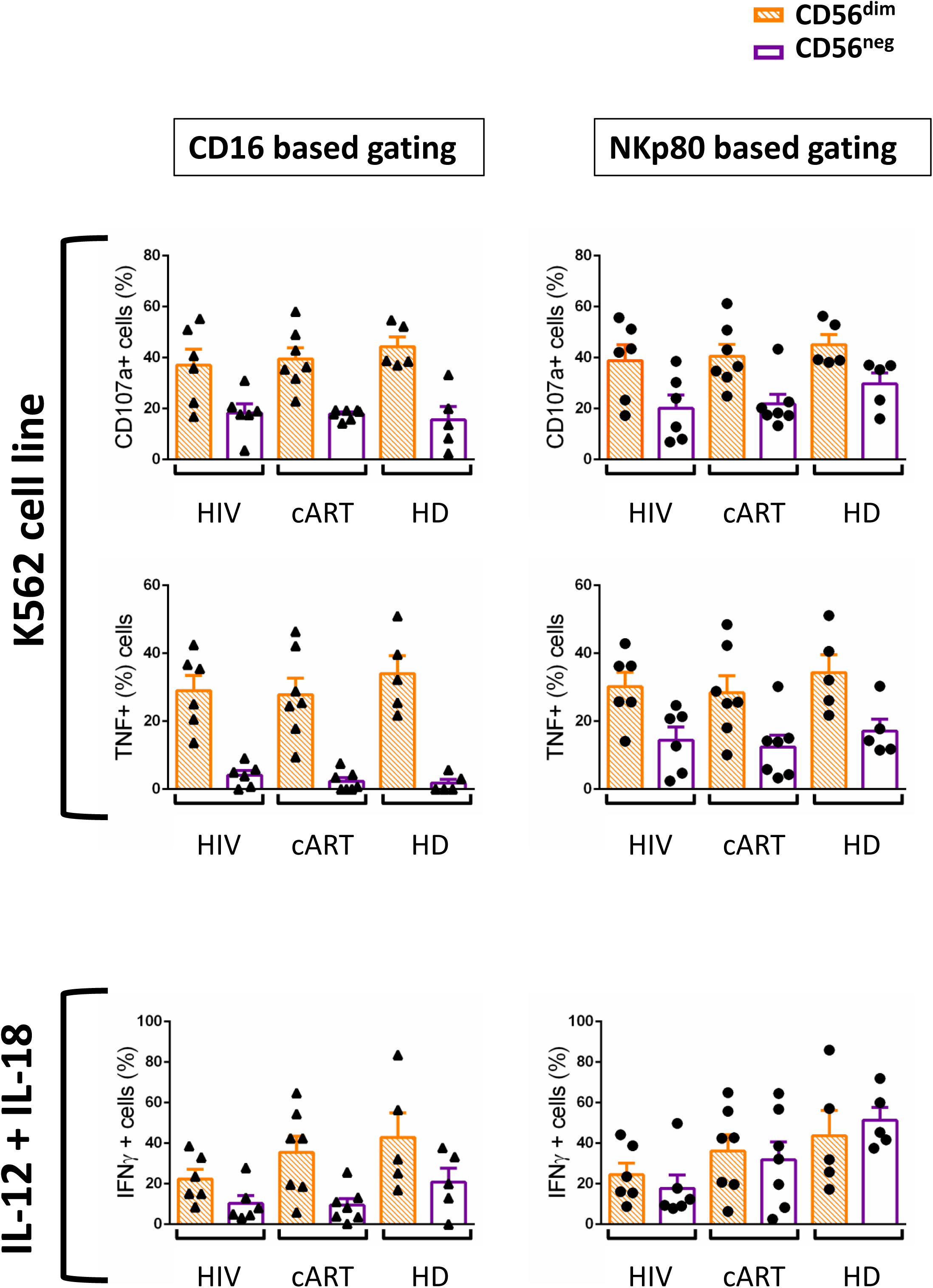
Degranulation (CD107a) and cytokine production by NK cells in response to the K562 cell line and IL-12+IL-18 cytokine stimulation. Bar graphs showing the percentage of CD56^dim^ and CD56^neg^ NK cells positive for CD107a and TNF after K562 cell line stimulation and IFNγ after IL-12+IL-18 cytokine stimulation from HIV-1 infected subjects (HIV), HIV-1 infected patients under cART (cART) and healthy donors (HD). The mean with the standard error of the mean (SEM) is represented. Each dot represents a donor.

**Supplementary Table 1.** Clinical data of untreated HIV-1 infected subjects, under cART HIV-1 infected patients and multiple myeloma patients.

|  | Untreated HIV-1 subjects |  | HIV-1 patients on cART |  | Multiple myeloma patients |  |
| --- | --- | --- | --- | --- | --- | --- |
|  | Median | Range (min-max) | Median | Range (min-max) | Median | Range (min-max) |
| **Sex** | Female: n=0<br>Male: n=9 | - | Female: n=1<br>Male: n=7 | - | Female: n=6<br>Male: n=3 | - |
| **Age (years)** | 39 | (32-59) | 49 | (34-56) | 63 | (53-74) |
| **cART (months)** | - | - | 13 | (10-27) | - | - |

## Appendix I: CoRIS Members

**Executive committee**

Santiago Moreno, Inma Jarrín, David Dalmau, Maria Luisa Navarro, Maria Isabel González, Federico Garcia, Eva Poveda, Jose Antonio Iribarren, Félix Gutiérrez, Rafael Rubio, Francesc Vidal, Juan Berenguer, Juan González, M Ángeles Muñoz-Fernández.

**Fieldwork data management and analysis**

Inmaculada Jarrin, Belén Alejos, Cristina Moreno, Carlos Iniesta, Luis Miguel Garcia Sousa, Nieves Sanz Perez, Marta Rava

**BioBanK HIV Hospital General Universitario Gregorio Marañón**

M Ángeles Muñoz-Fernández, Irene Consuegra Fernández

**Hospital General Universitario de Alicante (Alicante)**

Esperanza Merino, Gema García, Irene Portilla, Iván Agea, Joaquín Portilla, José Sánchez-Payá., Juan Carlos Rodríguez, Lina Gimeno, Livia Giner, Marcos Díez, Melissa Carreres, Sergio Reus, Vicente Boix, Diego Torrús

**Hospital Universitario Central de Asturias (Oviedo)**

Víctor Asensi, Eulalia Valle, María Eugenia Rivas Carmenado, Tomas Suarez-Zarracina Secades, Laura Pérez Is

**Hospital Universitario 12 de Octubre (Madrid)**

Rafael Rubio, Federico Pulido, Otilia Bisbal, Asunción Hernando, Lourdes Domínguez, David Rial Crestelo, Laura Bermejo, Mireia Santacreu

**Hospital Universitario de Donostia (Donostia-San Sebastián)**

José Antonio Iribarren, Julio Arrizabalaga, María José Aramburu, Xabier Camino, Francisco Rodríguez-Arrondo, Miguel Ángel von Wichmann, Lidia Pascual Tomé, Miguel Ángel Goenaga, Mª Jesús Bustinduy, Harkaitz Azkune, Maialen Ibarguren, Aitziber Lizardi, Xabier Kortajarena., Mª Pilar Carmona Oyaga, Maitane Umerez Igartua

**Hospital General Universitario De Elche (Elche)**

Félix Gutiérrez, Mar Masiá, Sergio Padilla, Catalina Robledano, Joan Gregori Colomé, Araceli Adsuar, Rafael Pascual, Marta Fernández, José Alberto García, Xavier Barber, Vanessa Agullo Re, Javier Garcia Abellan, Reyes Pascual Pérez, María Roca

**Hospital General Universitario Gregorio Marañón (Madrid)**

Juan Berenguer, Juan Carlos López Bernaldo de Quirós, Isabel Gutiérrez, Margarita Ramírez, Belén Padilla, Paloma Gijón, Teresa Aldamiz-Echevarría, Francisco Tejerina, Francisco José Parras, Pascual Balsalobre, Cristina Diez, Leire Pérez Latorre., Chiara Fanciulli

**Hospital Universitari de Tarragona Joan XXIII (Tarragona)**

Francesc Vidal, Joaquín Peraire, Consuelo Viladés, Sergio Veloso, Montserrat Vargas, Montserrat Olona, Anna Rull, Esther Rodríguez-Gallego, Verónica Alba., Alfonso Javier Castellanos, Miguel López-Dupla

**Hospital Universitario y Politécnico de La Fe (Valencia)**

Marta Montero Alonso, José López Aldeguer, Marino Blanes Juliá, María Tasias Pitarch, Iván Castro Hernández, Eva Calabuig Muñoz, Sandra Cuéllar Tovar, Miguel Salavert Lletí, Juan Fernández Navarro.

**Hospital Universitario La Paz/IdiPAZ**

Juan González-Garcia, Francisco Arnalich, José Ramón Arribas, Jose Ignacio Bernardino de la Serna, Juan Miguel Castro, Ana Delgado Hierro, Luis Escosa, Pedro Herranz, Víctor Hontañón, Silvia García-Bujalance, Milagros García López-Hortelano, Alicia González-Baeza, Maria Luz Martín-Carbonero, Mario Mayoral, Maria Jose Mellado, Rafael Esteban Micán, Rocio Montejano, María Luisa Montes, Victoria Moreno, Ignacio Pérez-Valero, Guadalupe Rúa Cebrián, Berta Rodés, Talia Sainz, Elena Sendagorta, Natalia Stella Alcáriz, Eulalia Valencia.

**Hospital Universitari MutuaTerrassa (Terrasa)**

David Dalmau, Angels Jaén, Montse Sanmartí, Mireia Cairó, Javier Martinez-Lacasa, Pablo Velli, Roser Font, Marina Martinez, Francesco Aiello

**Hospital Universitario de La Princesa (Madrid)**

Ignacio de los Santos, Jesus Sanz Sanz, Ana Salas Aparicio, Cristina Sarria Cepeda, Lucio Garcia-Fraile Fraile, Enrique Martín Gayo.

**Hospital Universitario Ramón y Cajal (Madrid)**

Santiago Moreno, José Luis Casado Osorio, Fernando Dronda Nuñez, Ana Moreno Zamora, Maria Jesús Pérez Elías, Carolina Gutiérrez, Nadia Madrid, Santos del Campo Terrón, Sergio Serrano Villar, Maria Jesús Vivancos Gallego, Javier Martínez Sanz, Usua Anxa Urroz, Tamara Velasco

**Hospital General Universitario Reina Sofía (Murcia)**

Enrique Bernal, Alfredo Cano Sanchez, Antonia Alcaraz García, Joaquín Bravo Urbieta, Angeles Muñoz Perez, Maria Jose Alcaraz, Maria del Carmen Villalba.

**Hospital Nuevo San Cecilio (Granada)**

Federico García, José Hernández Quero, Leopoldo Muñoz Medina, Marta Alvarez, Natalia Chueca, David Vinuesa García , Clara Martinez-Montes., Carlos Guerrero Beltran, Adolfo de Salazar Gonzalerz, Ana Fuentes Lopez

**Centro Sanitario Sandoval (Madrid)**

Montserrat Raposo Utrilla, Jorge Del Romero, Carmen Rodríguez, Teresa Puerta, Juan Carlos Carrió, Mar Vera, Juan Ballesteros, Oskar Ayerdi.

**Hospital Universitario Son Espases (Palma de Mallorca)**

Melchor Riera, María Peñaranda, Mª Angels Ribas, Antoni A Campins, Carmen Vidal, Francisco Fanjul, Javier Murillas, Francisco Homar., Helem H Vilchez, Maria Luisa Martin, Antoni Payeras.

**Hospital Universitario Virgen de la Victoria (Málaga)**

Jesús Santos, Crisitina Gómez Ayerbe, Isabel Viciana, Rosario Palacios, Carmen Pérez López, Carmen Maria Gonzalez-Domenec

**Hospital Universitario Virgen del Rocío (Sevilla)**

Pompeyo Viciana, Nuria Espinosa, Luis Fernando López-Cortés.

**Hospital Universitario de Bellvitge (Hospitalet de Llobregat)**

Daniel Podzamczer, Arkaitz Imaz, Juan Tiraboschi, Ana Silva, María Saumoy, Paula Prieto

**Hospital Costa del Sol (Marbella)**

Julián Olalla Sierra, Javier Pérez Stachowski., Alfonso del Arco, Javier de la Torre, José Luis Prada, José María García de Lomas Guerrero

**Hospital General Universitario Santa Lucía (Cartagena)**

Onofre Juan Martínez, Francisco Jesús Vera, Lorena Martínez, Josefina García, Begoña Alcaraz, Amaya Jimeno.

**Complejo Hospitalario Universitario a Coruña (Chuac) (A Coruña)**

Angeles Castro Iglesias, Berta Pernas Souto, Alvaro Mena de Cea.

**Hospital Universitario Virgen de la Arrixaca (El Palmar)**

Carlos Galera, Helena Albendin, Aurora Pérez, Asunción Iborra, Antonio Moreno, Maria Angustias Merlos, Asunción Vidal, Marisa Meca

**Hospital Universitario Infanta Sofia (San Sebastian de los Reyes)**

Inés Suárez-García, Eduardo Malmierca, Patricia González-Ruano, Dolores Martín Rodrigo, Mª Pilar Ruiz Seco.

**Hospital Universitario Príncipe de Asturias (Alcalá de Henares)**

José Sanz Moreno, Alberto Arranz Caso, Cristina Hernández Gutiérrez, María Novella Mena.

**Hospital Clínico Universitario de Valencia (València)**

María José Galindo Puerto, Ramón Fernando Vilalta, Ana Ferrer Ribera.

**Hospital Reina Sofía (Córdoba)**

Antonio Rivero Román, Antonio Rivero Juárez, Pedro López López, Isabel Machuca Sánchez, Mario Frias Casas, Angela Camacho Espejo

**Hospital Universitario Severo Ochoa (Leganés)**

Miguel Cervero Jiménez, Rafael Torres Perea

**Nuestra Señora de Valme (Sevilla)**

Juan A Pineda, Pilar Rincón Mayo, Juan Macias Sanchez, Nicolas Merchante Gutierrez, Luis Miguel Real, Anais Corma Gomez, Marta Fernandez Fuertes, Alejandro Gonzalez-Serna

